# Glyoxalase (GLO1) inhibition or genetic overexpression does not alter ethanol’s locomotor effects: implications for GLO1 as a therapeutic target in alcohol use disorders

**DOI:** 10.1101/239624

**Authors:** Amanda M Barkley-Levenson, Frances A Lagarda, Abraham A Palmer

**Author notes:** Corresponding author information: **Corresponding Author:** Abraham A. Palmer; Department of Psychiatry, University of California San Diego, La Jolla, CA 92093, USA. Voice.

## Abstract

**Background:** Glyoxalase 1 (**GLO1**) is an enzyme that metabolizes methylglyoxal (**MG**), which is a competitive partial agonist at GABA_A_ receptors. Inhibition of GLO1 increases concentrations of MG in the brain and decreases binge-like ethanol drinking. The present study assessed whether inhibition of GLO1, or genetic over expression of *Glo1*, would also alter the locomotor effects of ethanol, which might explain reduced ethanol consumption following GLO1 inhibition. We used the prototypical GABA_A_ receptor agonist muscimol as a positive control.

**Methods:** Male C57BL/6J mice were pretreated with aeither the GLO1 inhibitor S-bromobenzylglutathione cyclopentyl diester (**pBBG**; 7.5 mg/kg; Experiment 1) or muscimol (0.75 mg/kg; Experiment 2), or their corresponding vehicle. We then determined whether locomotor response to a range of ethanol doses (0, 0.5, 1.0, 1.5, 2.0, and 2.5) was altered by either pBBG or muscimol pretreatment. We also examined the locomotor response to a range of ethanol doses in FVB/NJ wild type and transgenic *Glo1* over expressing mice (Experiment 3). Anxiety-like behavior (time spent in the center of the open field) was assessed in all three experiments.

**Results:** The ethanol dose-response curve was not altered by pretreatment with pBBG or by transgenic overexpression of *Glo1*. In contrast, muscimol blunted locomotor stimulation at low ethanol doses, and potentiated locomotor sedation at higher ethanol doses. No drug or genotype differences were seen in anxiety-like behavior after ethanol treatment.

**Conclusions:** The dose of pBBG used in this study is within the effective range shown previously to reduce ethanol drinking. Glo1 overexpression has been previously shown to increase ethanol drinking. However, neither manipulation altered the dose response curve for ethanol’s locomotor effects, whereas muscimol appeared to enhance the locomotor sedative effects of ethanol. The present data demonstrate that reduced ethanol drinking caused by GLO1 inhibition is not due to potentiation of ethanol’s stimulant or depressant effects.

## Introduction

Alcohol use disorders (**AUD**) affect more than 15 million people in the United States alone (SAMHSA, 2015) and impose a cost of $249 billion annually (Sacks et al., 2015). Currently, there are only 3 FDA-approved medications to treat AUD, and these treatments are only affective in a subset of individuals (e.g. Garbutt, 2009; Maisel et al., 2013; Synder & Bowers, 2008). Consequently, there is a significant need for novel therapeutic targets for AUD.

One possible target is glyoxalase 1 (GLO1; McMurray et al., 2017a). GLO1 is a ubiquitously expressed enzyme that metabolizes the glycolytic byproduct methylglyoxal (**MG**). We have previously shown that MG acts as a competitive partial agonist at GABAA receptors (Distler et al., 2012). Pharmacological inhibition of GLO1 results in elevated MG levels in the brain, thus increasing GABAergic tone. Inhibition of GLO1 decreased binge-like ethanol drinking in mice during a drinking in the dark paradigm, as well as various anxiety- and depression-like behaviors (Distler et al., 2012, McMurray et al., 2017a, 2017b). In addition, transgenic overexpression of *Glo1* increased ethanol drinking (McMurray et al., 2017a).

Substantial previous work with various GABAergic agonists and positive allosteric modulators indicates that compounds that increase GABAergic activity can produce additive effects when administered with ethanol. Specifically, it has been well-established that enhanced locomotor sedation and ataxia is observed when ethanol and GABAergic agonists or positive modulators are co-administered (e.g. Tran et al., 2017; Vanover et al., 1999; Saeed Dar, 2006; Holstein et al., 2009; Dudek & Phillips, 1989). Additive locomotor effects with ethanol could potentially explain a compound’s ability to reduce ethanol consumption due to an increase in competing behaviors (e.g. locomotor activation) or through increased sedative effects that limit consumption. In humans, locomotor stimulatory effects of ethanol are associated with euphoria and drug-liking, whereas sedation is associated with more negative/aversive drug responses (Fridberg et al., 2017; King et al., 2011). Thus, measurements of ethanol-induced changes in locomotor behavior may also provide some insight into sensitivity to ethanol’s subjective effects.

We have shown previously that GLO1 inhibition does not alter the duration of ethanol-induced loss of righting reflex or ataxia as assessed by footslips on a balance beam (McMurray et al., 2017a). However, these studies only considered one ethanol dose and did not measure locomotor activity. Consequently, we sought to explore whether increased GLO1 inhibition (and subsequently increased GABAA receptor activation) produced additive effects on locomotor behavior when co-administered with a range of ethanol doses, thereby providing a possible explanation for the previously observed reductions in ethanol consumption. To test this hypothesis, we assessed the effects of GLO1 inhibition on the locomotor response to different doses of ethanol. As a positive control, we also assessed the effects of the non-selective GABA_A_ agonist muscimol, which has been previously shown to potentiate the sedative effects of ethanol (Holstein et al., 2004). We selected doses of pBBG and Muscimol that had been previously shown to inhibit ethanol drinking (McMurray et al., 2017a; Quoilin & Boehm, 2016). To further explore whether GLO1 alterations affect the locomotor response to ethanol, we also assessed the effect of *Glo1* overexpression on the locomotor response to ethanol. Transgenic mice overexpressing *Glo1* on an FVB/NJ (FVB) genetic background have been previously shown to have greater binge-like ethanol intake than wild type littermates (McMurray et al., 2017a) and have lower brain MG concentrations (Distler et al., 2012). If MG and ethanol act additively on ethanol locomotor response, then we would expect *Glo1* transgenic mice to be less sensitive to the locomotor effects of ethanol. Additionally, because Glo1 genetic and pharmacological manipulations have been shown previously to alter anxiety-like behavior (i.e. *Glo1* overexpression increases anxiety-like behavior and Glo1 inhibition decreases it; Distler et al. 2012), we also measured anxiety-like behavior in these studies to assess potential treatment interactions with ethanol anxiolytic response.

## Materials and Methods

### Animals and husbandry

For experiments 1 and 2, 8 week old male C57BL/6J (B6) mice were ordered from the Jackson Laboratory (Bar Harbor, ME). All mice were between 65 and 74 days of age at the time of testing. For experiment 3, male and female mice heterozygous for *Glo1* overexpression on a FVB background and wild type littermates were bred in house. Generation of the *Glo1* transgenic mice by insertion of a BAC transgene has been previously described (Distler et al., 2012). Mice in experiment 3 were between 57 and 139 days of age at the time of testing. For all experiments, mice were housed 2-5 per cage on cob bedding and food (Envigo 8604, Indianapolis, IN) and water were provided *ad libitum*. Mice were maintained on a 12hr/12hr light/dark cycle with lights on at 06:00. Behavioral testing was conducted during the light phase. All procedures were approved by the local Institutional Animal Care and Use Committees and were conducted in accordance with the NIH Guidelines for the Care and Use of Laboratory Animals.

### Open field testing and apparatus

Mice were moved into the testing room and allowed to acclimate for at least 30 min prior to testing. Locomotor behavior was measured using an open field test, which has been described previously (McMurray et al., 2016). Briefly, mice were placed in the center of a square chamber (43 ×43 × 33 cm) (Accusan, Columbus, OH) with dim overhead lighting inside of sound- and light-attenuating boxes and allowed to freely explore for 30 min. A grid of infrared detection beams in each chamber and Versamax software was used to track animal location and locomotor activity (distance traveled) during the test. Although not directly related to the main hypothesis of these experiments, we also chose to record time spent in the center zone (26 × 26 cm) as a measure of anxiety-like behavior to assess potential interactions between the Glo1 manipulations and ethanol anxiolytic response, since both Glo1 inhibition and ethanol can produce anxiolytic effects. The chambers were wiped down with isopropyl alcohol between each animal to eliminate odors.

### Drugs

The GLO1 inhibitor S-bromobenzylglutathione cyclopentyl diester (pBBG) was synthesized in the laboratory of Alexander Arnold at the University of Wisconsin Milwaukee, as previously described (McMurray et al., 2017a). The pBBG dose (7.5 mg/kg) was chosen because it is in the range shown to reduce binge-like ethanol drinking (6.25-12.5 mg/kg, McMurray et al., 2017a). pBBG was dissolved in vehicle (8% DMSO/18% tween80/saline) and administered i.p. (injection volume 0.01 ml/g). The pBBG pretreatment was given 2 h before testing, consistent with previous studies demonstrating pBBG effects on ethanol drinking, anxiety-like, and depression-like behaviors (McMurray et al., 2017a, 2017b; Distler et al., 2012)., Muscimol was purchased from Sigma-Aldrich (St. Louis, MO, USA). The muscimol dose (0.75 mg/kg) was chosen because it has been shown previously to reduce ethanol drinking (Quoilin & Boehm, 2016) but was not expected to produce locomotor effects in the absence of co-administration of ethanol (Holstein et al., 2009; Shen et al., 1998). Muscimol was dissolved in 0.9% saline and administered i.p. (injection volume 0.01 ml/g). All mice in a squad (8 mice/squad) were first injected with muscimol and then all mice were injected with ethanol. Time between muscimol and ethanol injections for each animal was approximately 4 minutes which is similar to previous studies (Holstein et al., 2009). In contrast, in experiment 1, pBBG was administered 2 h prior to ethanol injection to allow for MG accumulation. Ethanol (Deacon Laboratories Inc., King of Prussia, PA) was dissolved in saline (20% v/v) and administered i.p. at a dose of 0, 0.5, 1, 1.5, 2, or 2.5 g/kg. This dose range was chosen to encompass both locomotor stimulatory and sedative effects.

### Experiment 1: pBBG effects on ethanol locomotor and anxiolytic response

144 mice were used (n=12/drug/ethanol dose) for this experiment. Mice were psuedorandomly assigned to a pretreatment group of either pBBG or vehicle (8% DMSO/18% tween80/saline), and an ethanol dose group (0 [saline], 0.5, 1, 1.5, 2, or 2.5 g/kg). Squads of eight mice were injected with their assigned pretreatment compound and returned to the home cage for 2 h. We have previously used the 2 h time point because it is believed to allow for the accumulation of behaviorally-relevant MG concentrations in the brain (Distler et al., 2012, 2013) and critically because it was used in the studies that showed inhibition of ethanol drinking following pBBG pretreatment (McMurray 2017a). Two hours later, mice were injected i.p. with their assigned ethanol dose and immediately placed in the activity chambers for 30 minutes.

### Experiment 2: muscimol effects on ethanol locomotor and anxiolytic response

142 mice were used (n=11-12/drug/ethanol dose) for this experiment. Mice were psuedorandomly assigned to a pretreatment group of either the GABA_A_ agonist muscimol or vehicle (saline) and an ethanol dose group (0 [saline], 0.5, 1, 1.5, 2, or 2.5 g/kg). Squads of eight mice were injected with their assigned pretreatment compound immediately before a second injection of ethanol, as described in experiment 1. Mice were then immediately placed in the activity chambers for 30 minutes.

### Experiment 3: ethanol locomotor and anxiolytic response in Glo1 transgenic mice

152 male and female Glo1 overexpressing mice and wild type littermates were used in this experiment (n=6-14/genotype/dose). Wild type FVB mice show locomotor stimulation at higher ethanol doses than B6 mice (Metten et al., 2004), so we extended the tested dose range to 3 g/kg to capture more of the dose-response curve. Mice were pseudorandomly assigned to an ethanol dose group (0 [saline], 0.5, 1, 1.5, 2, 2.5, or 3 g/kg). Squads of eight mice were injected with ethanol and immediately placed in the activity chambers for 30 minutes.

### Statistical analyses

Locomotor activity was analyzed as total distance traveled in 30 min, and as a total of 6 bins of 5 min each in order to assess drug and time course interactions (experiments 1 and 2). Statistical analyses were performed in SPSS (version 24; IBM Corp, Armonk, NY). Total distance data were analyzed by two-way analysis of variance (ANOVA) with the between-groups factors of drug (pBBG vs vehicle or muscimol vs vehicle) and ethanol dose (0, 0.5, 1.0, 1.5, 2.0, and 2.5 g/kg) for experiments 1 and 2, and genotype (transgenic vs wild type) and ethanol dose (0, 0.5, 1.0, 1.5, 2.0, 2.5, and 3.0 g/kg) for experiment 3. Significant interactions were followed up by one-way ANOVAs; post hoc Dunnett tests were also used to compare ethanol doses to the saline treated group. Time course data were analyzed using a two-way mixed model ANOVA with between-groups factors of drug (pBBG vs vehicle or muscimol vs vehicle) and ethanol dose (0, 0.5, 1.0, 1.5, 2.0, and 2.5 g/kg), and with time as a within-subjects repeated measure (6 total bins of 5 min each). A Huyhn-Feldt correction was used for repeated measures ANOVA where appropriate. Significant interactions were followed up by lower order ANOVAs. Center times were converted to a percent of total test time and were analyzed using two-way analysis of variance (ANOVA). Significant interactions were followed up by one-way ANOVAs and post hoc Dunnett tests were also used to compare ethanol doses to the saline treated group.Significance was set at α=0.05 for all tests.

We could not compare pBBG to muscimol because experiments 1 and 2 were performed separately and because both pretreatment time (2 h versus 0 min) and the vehicle solution were different.

## Results

### Experiment 1: pBBG effects on ethanol locomotor response

Figure 1 shows total distance traveled after each ethanol dose for pBBG- and vehicle-treated mice. Statistical analyses showed a main effect of ethanol dose (*F*_5,143_=31.11, p<0.001), but no main effect of drug or a significant drug x ethanol dose interaction. Dunnett post hoc tests comparing each ethanol dose to saline (collapsed on drug) showed that the 1 and 1.5 g/kg doses of ethanol produced a significant locomotor stimulatory effect (p=0.005 and p=0.021, respectively). In contrast, the 2 and 2.5 g/kg doses of ethanol produced significant locomotor sedation compared to saline (p<0.001 for both).

**Figure 1:**
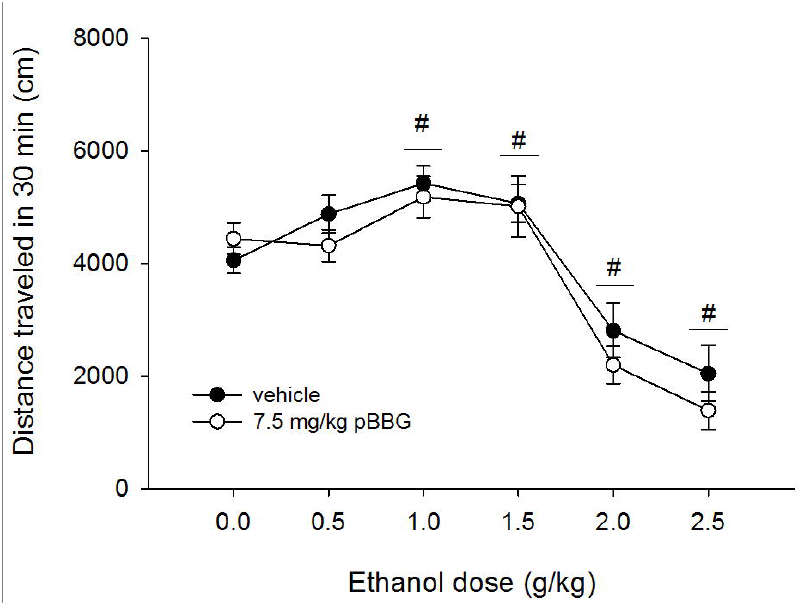
Total distance traveled in 30 min for each dose of ethanol (EtOH) in mice treated with 7.5 mg/kg of the GLO1 inhibitor pBBG or vehicle. Significant results of the post-hoc tests for the main effect of ethanol dose (*F*_5,143_=31.11, p<0.001) are noted, where # indicates the entire dose group (collapsed on inhibitor treatment) is significantly different from 0 g/kg (saline) ethanol group.

Figure 2 (a-f) shows the time course of locomotor activity for pBBG- and vehicle-treated animals at each ethanol dose. Mixed-model ANOVA showed a main effect of time (*F*_3.19,421.41_ =292.24, p<0.001) and a significant time x ethanol dose interaction (*F*_15.9,421.41_=7.5, p<0.001). There was no significant time x drug x ethanol dose interaction or time x drug interaction. Because of our *a priori* interest in potential differences in pretreatment effects at sedative vs stimulatory ethanol doses, we chose to examine each ethanol dose individually. There was no main effect of drug and no significant time x drug interaction found for any dose of ethanol (p≥0.29 for all), indicating that pBBG treatment did not alter ethanol locomotor response at any time point or ethanol dose (see Figure 2).

**Figure 2:**
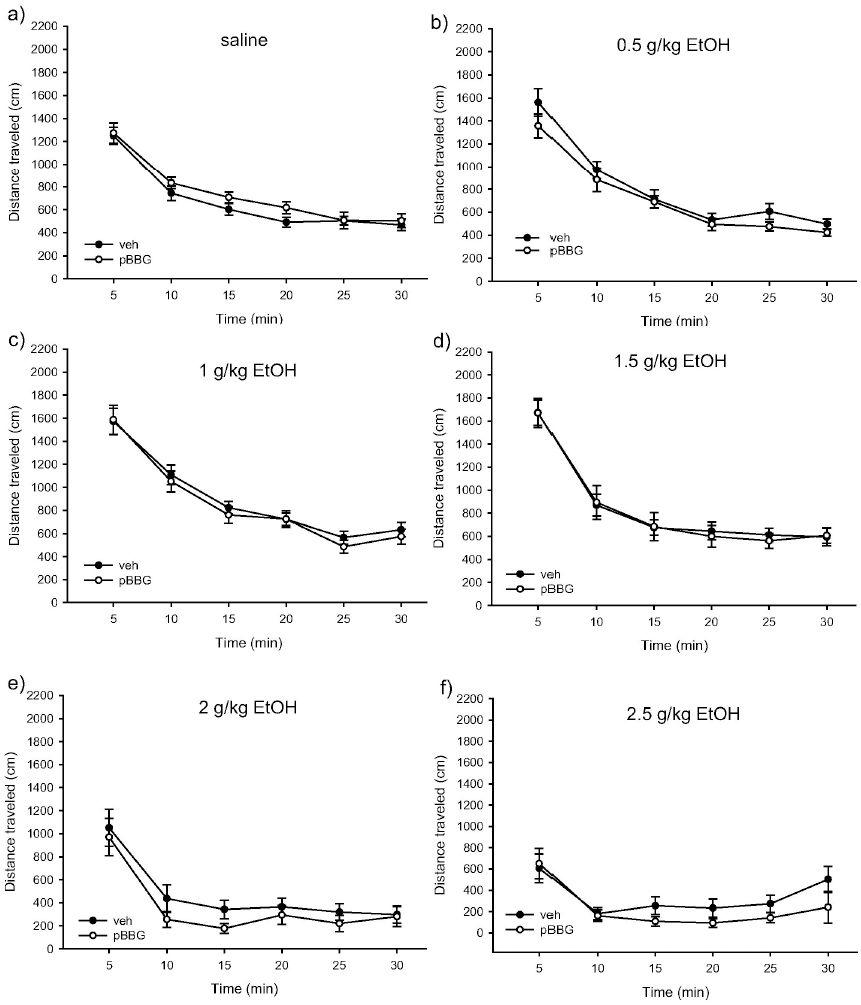
Time courses of locomotor activity by ethanol dose for pBBG- and vehicle-treated groups. No significant main effects or interactions were seen for the drug groups.

### Experiment 1: pBBG effects on ethanol anxiolytic response

Figure 3 shows the percent of total time spent in the center region after each ethanol dose for pBBG- and vehicle-treated mice. Time course center time data and analyses are shown in supplemental figure S1. Statistical analyses showed a main effect of ethanol dose (*F*_5,144_=10, p<0.001) and a significant ethanol dose x drug interaction (*F*_5,144_=2.53, p=0.032). Follow up ANOVAs showed a main effect of drug only for the 0 g/kg ethanol dose group (saline control), where pBBG-treated mice spent significantly more time in the center than the vehicle-treated mice (*F*_1,24_=11.17, p=0.003).

**Figure 3:**
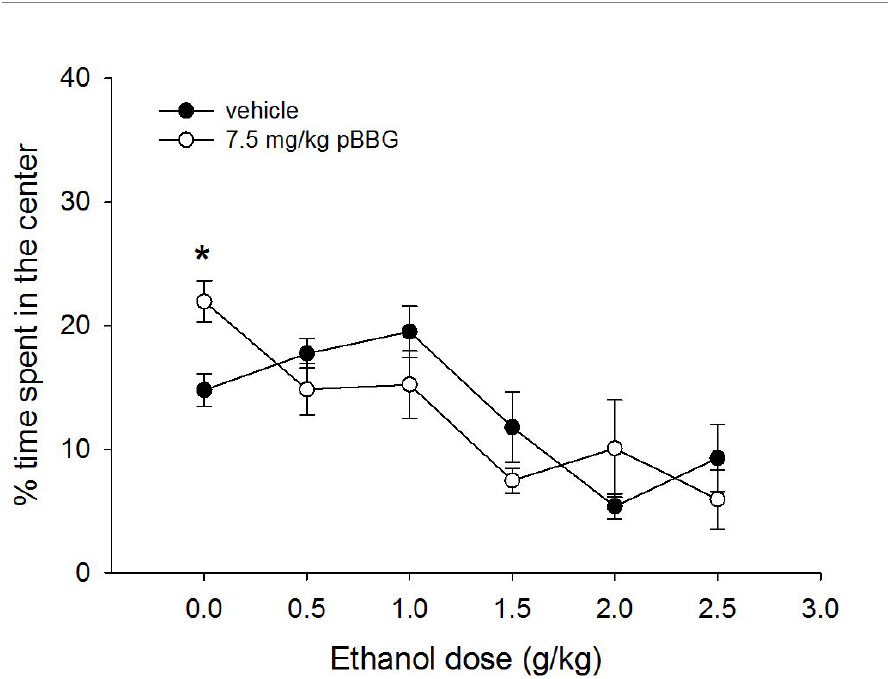
Percent of total time spent in the center of the open field apparatus during 30 min test for each dose of ethanol in mice treated with 7.5 mg/kg of the GLO1 inhibitor pBBG or vehicle. ^*^ indicates significant difference between pBBG and vehicle groups.

### Experiment 2: muscimol effects on ethanol locomotor response

Figure 4 shows total distance traveled after each ethanol dose for muscimol- and vehicle-treated mice. Statistical analyses showed a main effect of drug (*F*_1,142_=7.76, p<0.001), a main effect of ethanol dose (*F*_5.142_=14.04, p<0.001) and a significant drug x ethanol dose interaction (*F*_5,142_=2.86, p<0.017). Follow up ANOVAs were used to assess the effect of ethanol dose within the muscimol and vehicle groups separately. Both the vehicle and muscimol groups showed a main effect of ethanol dose (*F*_1,71_≥4.9, p≤0.001). However, we observed significant locomotor stimulation at the 1 g/kg dose in the vehicle group (1 g/kg >saline, p=0.015) that was not present in the muscimol-treated animals (p=0.23). Furthermore, the 2.5 g/kg ethanol dose produced a sedative effect in the muscimol treated group (p<0.001) but not in the vehicle treated group (p=0.83).

**Figure 4:**
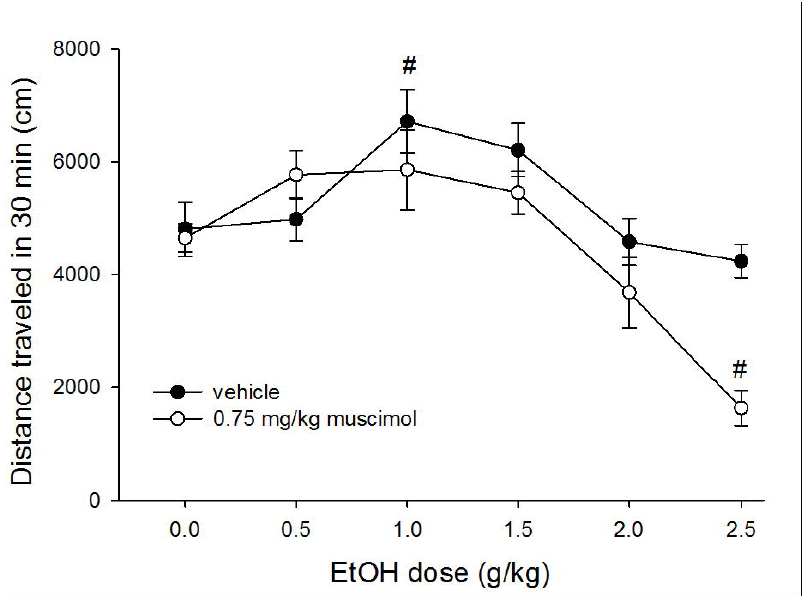
Total distance traveled in 30 min for each dose of ethanol in mice treated with 0.75 mg/kg muscimol or vehicle. # indicates statistically significant difference from the 0 g/kg (saline) group of the same drug treatment.

Figure 5 (a-f) shows the time course of locomotor activity for muscimol- and vehicle-treated animals at each ethanol dose. Mixed-model ANOVA showed a main effect of time (*F*_3.74_, 486.48=275.44, p<0.001), a significant time x drug interaction (*F*_3.74,48.48_=6.03, p<0.001), and a significant time x ethanol dose interaction (*F*_1.76,48.48_=2.64, p<0.001). There was also a trend toward a significant 3-way interaction of time x drug x ethanol dose (*F*_18.71,486.48_=1.53, p=0.073). Due to our *a priori* interest in individual ethanol dose effects (sedative vs. stimulatory), we chose to follow up this suggestive 3-way interaction by examining the effects of time and muscimol within each ethanol dose group. The 0 g/kg (saline), 0.5 g/kg, and 2 g/kg ethanol dose groups showed main effects of time (*F*_2.84-5,2.45-105_≥62.45, p<0.001), but no time x drug interactions and no main effects of drug. The 1 g/kg ethanol dose group also showed a main effect of time (*F*_3.53,77.68_=62.34, p<0.001) and no main effect of drug, though there was a trend toward a significant time x drug interaction (*F*_2.89,77.68_=2.59, p=0.063). The 1.5 g/kg ethanol dose group showed a main effect of time (*F*_4.37,96.12_=53.97, p<0.001), no main effect of drug, and a significant time x drug interaction (*F*_4.37,6.12_=6.24, p<0.001). Follow up ANOVAs for each time bin found that muscimol treated mice had significantly less locomotor activity during the 10 and 15 min time bins than vehicle treated mice (**Figure 3d**; main effect of drug, *F*_1,24_=12.61, p=0.002 and *F*_1,24_=8.21, p=0.009, respectively). The 2.5 g/kg ethanol dose group also showed a trend toward a significant time x drug interaction (*F*_3.82,83.82_=2.5, p=0.051), and main effects of both time and drug (*F*_2.81,83.82_=30.06, p<0.001 and *F*_1,22_=36.88, p<0.001, respectively), with muscimol treated animals showing significantly more locomotor sedation than vehicle treated animals for all but one time point (**Figure 3f**; *F*_1,24_≥10.55, p≤0.004).

**Figure 5:**
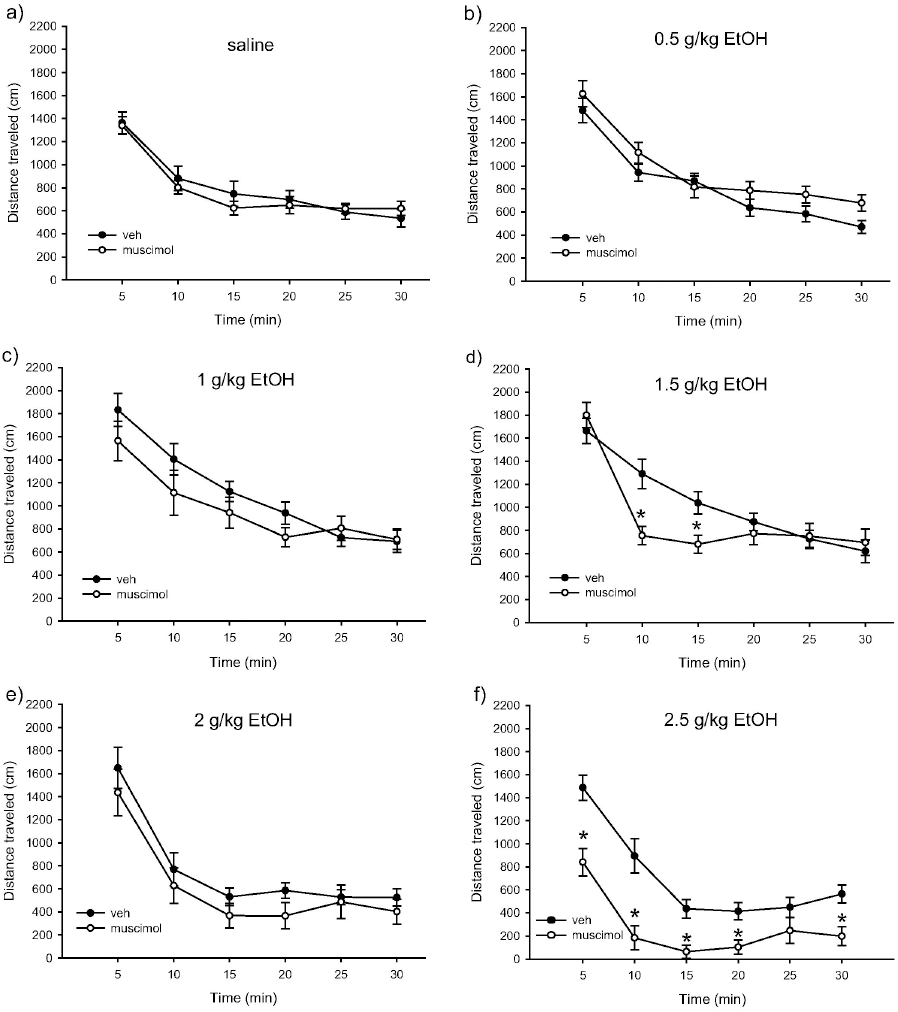
Time courses of locomotor activity by ethanol dose for muscimol- and vehicle-treated groups. ^*^ indicates statistically significant difference from the vehicle treated group for a given time bin.

### Experiment 2: muscimol effects on ethanol anxiolytic response

Figure 6 shows the percent of total time spent in the center region after each ethanol dose for muscimol-and vehicle-treated mice. Time course center time data and analyses are shown in supplemental figure S2. One muscimol-treated animal in the 2 g/kg ethanol group was excluded as an outlier (center time > 3 standard deviations from the mean). Statistical analyses showed main effects of ethanol dose and drug (*F*_5,141_=5.77, p<0.001 and *F*_1,141_ =4.7, p=0.032, respectively), and a significant ethanol dose x drug interaction (*F*_5,141_=3.06, p=0.012). Follow up ANOVAs showed a main effect of drug for the 2.5 g/kg ethanol group (*F*_1,24_=4.77, p=0.04), and a trend toward a main effect for the 0.5 g/kg ethanol group (*F*_1,23_=3.46, p=0.077). For the 2.5 g/kg ethanol group, the muscimol-treated mice spent more time in the center than the vehicle-treated mice. However, caution must be used when interpreting this difference due to the profound locomotor sedation seen in the muscimol-treated mice in this dose group; while there were no statistical outliers, 8 out of 12 muscimol-treated mice in the 2.5 g/kg ethanol group spent at least 10 consecutive minutes with no locomotor activity, suggesting significant sedation that would interfere with the expression of anxiety-like behavior in this test.

**Figure 6:**
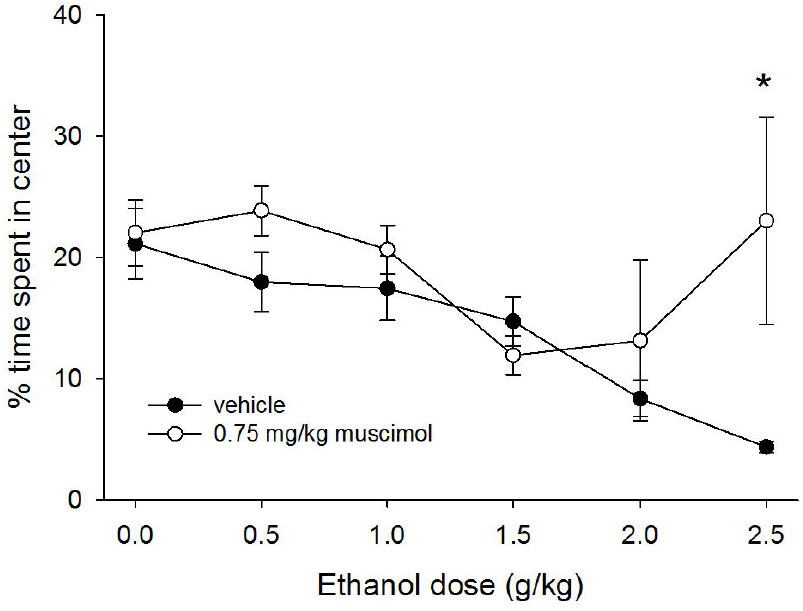
Percent of total time spent in the center of the open field apparatus during 30 min test for each dose of ethanol in mice treated with 0.75 mg/kg muscimol or vehicle. ^*^ indicates statistically significant difference between the muscimol and vehicle groups.

### Experiment 3: ethanol locomotor response in Glo1 transgenic mice

Figure 7 shows total distance traveled after each ethanol dose for *Glo1* overexpressing mice and wild type littermates. Statistical analyses showed main effects of sex (females>males, *F*_1,152_=7.52, p=0.007) and ethanol dose (*F*_6,152_=11.77, p<0.001), but no main effect of genotype and no significant interactions. Because there were no significant interactions between sex and other variables, all data are presented collapsed on sex. Dunnett post hoc tests showed that there was significant locomotor stimulation at the 1.5, 2, and 2.5 g/kg ethanol doses as compared to saline (p≤0.003 for all).

**Figure 7:**
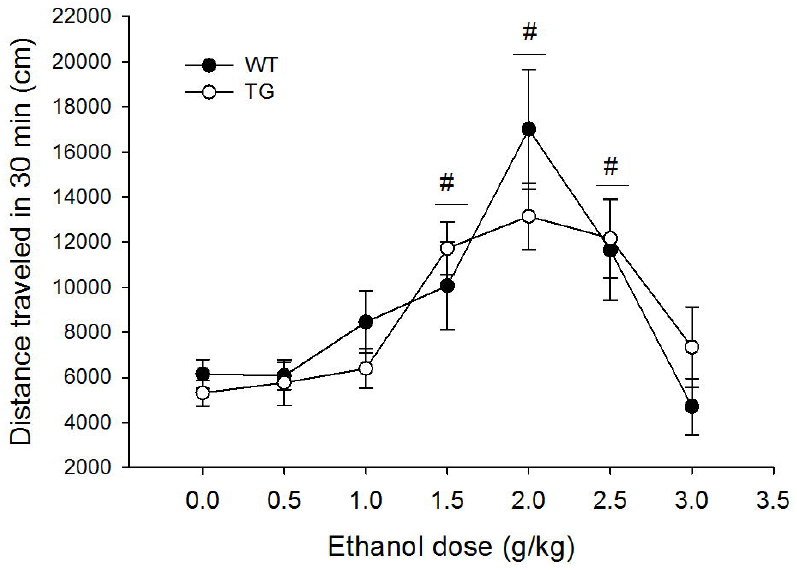
Total distance traveled in 30 min for each dose of ethanol in wild type (WT) and transgenic mice heterozygous for the *Glo1* over expression (**TG),** collapsed on sex. Significant results of the post-hoc tests for the main effect of ethanol dose (*F*_6,152_=11.77, p<0.001) are noted, where # indicates the entire dose group (collapsed on genotype) is significantly different from 0 g/kg (saline) ethanol group.

### Experiment 3: ethanol anxiolytic response in Glo1 transgenic mice

Figure 8 shows percent of total time spent in the center after each ethanol dose for *Glo1* overexpressing mice and wild type littermates. Time course center time data and analyses are shown in supplemental figure S3. Statistical analyses showed a main effect of ethanol dose (*F*_6,152_=2.80, p=0.013) but no significant interactions. Dunnett post hoc tests showed a significant anxiolytic effect of ethanol at the 3 g/kg dose (p=0.037).

**Figure 8:**
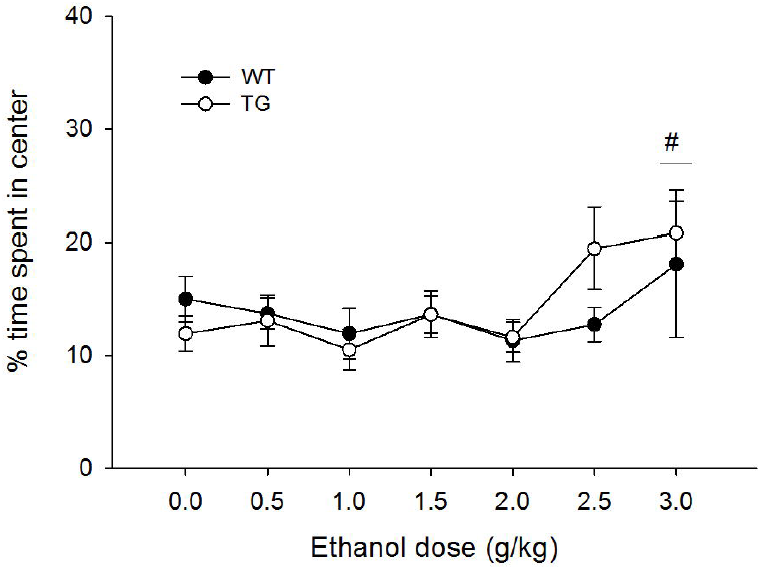
Percent of total time spent in the center of the open field apparatus during 30 min test for each dose of ethanol in wild type (WT) and transgenic mice heterozygous for the *Glo1* over expression (TG), collapsed on sex. Significant results of the post-hoc tests for the main effect of ethanol dose (*F*_6,151_= 3.22, p=0.006) are noted, where # indicates the entire dose group (collapsed on genotype) is significantly different from 0 g/kg (saline) ethanol group.

## Discussion

We found that a dose of the GLO1 inhibitor pBBG (7.5 mg/kg) that is within the effective range for reducing ethanol drinking did not alter the ethanol locomotor dose-response curve and had no effect on locomotor behavior when administered alone. In contrast, a dose of muscimol that has also been shown to reduce ethanol consumption did produce significant changes in the locomotor response to ethanol (attenuated stimulatory response and potentiated sedative response). Like pBBG, the dose of muscimol we used did not produce locomotor effects in the absence of ethanol. These findings suggest that unlike the GABAA receptor agonist muscimol, pBBG does not alter either the locomotor stimulant or depressant effects of ethanol. Further supporting these findings, *Glo1* transgenic mice showed locomotor responses that were similar to wild type littermates following a range of ethanol doses. These mice were of a different genetic background (FVB vs B6), so direct comparisons with the pharmacological experiments are somewhat complicated. However, the similar pattern of results between Experiments 1 and 3 (i.e. no effect of Glo1 inhibitor treatment or genetic overexpression of Glo1) does suggest that these findings generalize across genotypes. This absence of an effect on of Glo1 manipulations on ethanol locomotor response, especially ethanol locomotor sedation, is surprising given the considerable literature showing that other GABAergic drugs act synergistically with ethanol to potentiate sedative effects (e.g. Tran et al., 2017; Vanover et al., 1999; Saeed Dar, 2006; Holstein et al., 2009; Dudek & Phillips, 1989).

Although both pBBG and muscimol modulate GABAergic signaling, there are a number of important differences between them which may explain the differences in their interaction with ethanol. MG is a competitive partial agonist at GABA_A_ receptors (Distler et al., 2012), whereas muscimol is a competitive full agonist at GABA_A_ receptors (Arnt et al., 1979; Kemp et al., 1986). In addition, MG and muscimol likely have different GABA_A_ receptor subtypes specificity. Another difference is that treatment with pBBG causes the accumulation of MG that is proportionate to local glycolytic activity, whereas muscimol concentrations are expected to be uniform throughout the brain. These differences may explain why GLO1 inhibition (and the resulting elevation of MG) does not produce the additive effect with ethanol that was observed following muscimol pretreatment. Previous studies have shown that neither pBBG treatment nor direct MG administration potentiated other effects of ethanol; specifically, 50 mg/kg MG co-administered with 1.25 g/kg ethanol did not produce an increase in ethanol-induced ataxia as compared to ethanol administered alone. Similarly, two doses of pBBG shown to reduce drinking (6.25 mg/kg and 12.5 mg/kg) did not alter either ethanol-induced ataxia or the duration of ethanol-induced loss of righting reflex (McMurray et al., 2017a). Blood ethanol concentrations were not measured in these experiments, but muscimol has not been reported to alter ethanol metabolism; pBBG effects on ethanol metabolism have not be directly tested, but mice with reduced GLO1 expression (which show a similar reduction in ethanol consumption as pBBG-treated mice) do not differ in ethanol metabolism (McMurray et al. 2015). We therefore do not believe that differences in ethanol metabolism are likely to explain the observed differences in effects between pBBG and muscimol. Another possible explanation for the differences between pBBG and muscimol is that pBBG was injected 2 h before ethanol treatment, whereas muscimol was injected immediately before ethanol injection. Although this procedural difference is somewhat confounding, we chose these pretreatment times because they match the conditions used to inhibit ethanol drinking and because they did not produce changes in locomotor activity in the absence of ethanol. However, the gradual buildup of MG following pBBG pretreatment vs. the rapid onset of muscimol’s effects could explain the different interactions with ethanol-induced locomotor behavior.

Locomotor activity appeared to be lower in experiment 1 than in experiment 2 across all ethanol doses, particularly in the vehicle groups. There were several procedural differences between the two studies that might explain these differences. First, the vehicle solution used in experiment 1 (DMSO/Tween80/saline) was different from the vehicle solution used in experiment 2 (normal saline). The concentrations of DMSO and Tween80 used in experiment 1 (8% and 18%, respectively) are well below concentrations that have been previously reported to reduce locomotor activity in mice (Castro et al., 1995), but it remains a formal possibility that the difference in vehicle solution caused the differences in locomotor response among vehicle treated animals. Additionally, as discussed above, mice in experiment 1 had a longer pretreatment period (2 hours) as compared to mice in experiment 2, which could also have contributed to differences in locomotor response to ethanol. Ethanol locomotor effects in B6 mice can be surprisingly variable. For example, a 2 g/kg dose of ethanol has been shown to produce locomotor stimulation (Lessov et al., 2001), sedation (Sharko & Hodge, 2008; Dudek et al., 1991; Phillips & Dudek, 1991), and no effect (Phillips et al., 1995; Melón & Boehm, 2011; Camarini & Hodge, 2004; Rose et al., 2013), and even 2.5 g/kg does not always produce locomotor sedation (e.g. Hilbert et al., 2013). Therefore although the differences in vehicle groups is somewhat perplexing, the most relevant comparisons are between each drug and its corresponding vehicle group, as all mice in the same experiment received comparable treatments. At all ethanol doses tested, the vehicle-treated animals still showed sufficient locomotor activity to see a potentiation of sedative effects had one been present. Therefore we do not believe that group differences are being masked by a floor effect and the apparent differences in locomotor activity between experiments 1and 2 are unlikely to undermine our primary conclusions.

Neither pBBG nor muscimol had any systematic effect on ethanol’s anxiolytic effects. Similarly,*Glo1* transgenic mice did not differ from wild type mice in ethanol anxiolytic response. However, interpretation of these findings should be made cautiously given there was little evidence of an anxiolytic effect of ethanol alone in these experiments and some doses of ethanol appeared to reduce time spent in the center. Anxiety and locomotor activity can be extremely difficult to untangle due to the activity-dependent nature of the anxiety variables used (e.g. movement into/out of an “anxiogenic” region of the apparatus) (Milner & Crabbe, 2008).Therefore, the observed differences in locomotor behavior across dose groups may have confounded the anxiety results. For example, an increase in center time was seen in the 2.5 g/kg treated muscimol group in experiment 2. Upon closer examination of the data, many mice were clearly sedated (no activity for ≥10 min) and thus spent more time in the center of the open field. Therefore even an apparent “anxiolytic” effect of ethanol is complicated by the locomotor sedative effects. Ethanol has been shown to be anxiolytic in B6 mice at doses within the range used here (e.g. Gulick & Gould, 2009; Boyce-Rustay et al., 2007), However, these studies have used assays optimized for assessing anxiety-like behavior such as the elevated plus maze, and to our knowledge this effect has not been seen in the open field test we used here. Consequently, there may be apparatus-specific effects in ethanol anxiolytic response that are not apparent in the present experiments. However, we did observe a significant anxiolytic effect of pBBG treatment at baseline (0 g/kg ethanol), consistent with previous findings, but at a much lower dose of pBBG than has been seen before (50 mg/kg; Distler et al., 2012).

Although the GLO1 inhibitor did not produce the classically observed synergistic effect on locomotor sedation observed with other GABAergic drugs, it is possible that there are other additive effects with ethanol. For example, other drugs that increase GABAergic activity (e.g. barbiturates, benzodiazepines, neuroactive steroids) substitute for ethanol in drug discrimination procedures (Kostowski & Bienkowski, 1999; Shannon et al., 2004). Thus, increased MG activation of GABA_A_ receptors due to GLO1 inhibition may still produce interoceptive effects that are similar to ethanol itself. In this case, we might expect that GLO1 inhibitors decrease ethanol consumption by acting additively with ethanol in such a way that mice do not need to consume as much to achieve the same interoceptive effects.

In summary, we have shown that a dose of pBBG within the range that has been previously shown to reduce binge-like ethanol drinking does not alter sensitivity to ethanol locomotor stimulation or sedation, despite its presumed GABAergic mechanism of action. This is in direct contrast to our findings with the GABA_A_ agonist muscimol, which attenuated the locomotor stimulatory response to ethanol, and potentiated the sedative effect. This result suggests that ethanol drinking is not reduced by pBBG because it interferes with consumption by potentiating the sedative effects. Instead, GLO1 inhibition may reduce drinking by altering the rewarding or aversive properties of ethanol, a possibility that we are currently investigating. Overall, the lack of potentiation of ethanol’s locomotor effects seen with GLO1 inhibition suggest a mechanism that is qualitatively distinct from direct GABA agonists and modulators, and provides further support for the therapeutic potential of targeting this system in AUD.

## Acknowledgments

AMB-L was supported by NIH-NIAAA grant F32AA025515. The authors have no conflicts of interest to report.

---

**Figure S1:** Time course of center time by ethanol dose for pBBG- and vehicle-treated groups. There was no significant 3-way interaction of time x ethanol dose x drug group interaction, but there was a main effect of time (*F*_3.67,483.29_=3.02, p=0.021) and a significant time x ethanol dose interaction (*F*_18.31,483.29_=4.48, p<0.001).

**Figure S2:** Time course of center time by ethanol dose for muscimol- and vehicle-treated groups. There was no significant 3-way interaction of time x ethanol dose x drug group interaction, but there was a main effect of time (*F*_3.85,500.12_=7.02, p<0.001) and significant time x ethanol (*F*_3.95,500.12_=4.48, p=0.002) and time x drug group interactions (*F*_17.21,500.12_=2.29, p=0.002).

**Figure S3:** Time course of center time by ethanol dose for Glo1 transgenic (TG) and wild type (WT) mice. There were no main effects and no significant interactions.

